# Determining zygosity with multiplex Kompetitive Allele-Specific PCR (mxKASP) genotyping

**DOI:** 10.1101/2024.10.25.620256

**Authors:** Manon C. de Visser, Willem R.M. Meilink, Anagnostis Theodoropoulos, Michael Fahrbach, Ben Wielstra

## Abstract

We introduce multiplex Kompetitive Allele-Specific PCR (mxKASP): a modification of ‘classical’ KASP genotyping that allows zygosity to be determined in diploid organisms. Rather than targeting a SNP associated with a single marker, mxKASP targets two non-homologous markers. We show proof of concept by applying mxKASP to the balanced lethal system in *Triturus* newts, in which individuals are known to possess either: (1) zero copies of the 1A version of chromosome 1 and two copies of the 1B version; (2) one copy of 1A and one copy of 1B; or (3) two copies of 1A and zero copies of 1B. mxKASP is successful in amplifying both a 1A and a 1B marker in a single reaction (if present), allowing the zygosity of individuals to be inferred. We independently confirm our mxKASP results with a multiplex PCR approach. We argue that mxKASP can be applied to rapidly and economically determine zygostity in diploid organisms, for a large number of samples at once.

## Introduction

In diploid organisms, a stretch of DNA, such as a gene or even an entire chromosome, is *hemizygous* if only a single copy is present in the genome. The enormous impact (hemi)zygosity can have on the phenotype is perhaps best showcased by *sex chromosomes*. In species with genetic sex determination, the heterogametic sex is hemizygous for the region that underpins sex, whereas the homogametic sex is homozygous (Bachtrog et al., 2014). Zygosity also varies in the case of *supergenes*, which are physically linked genes that are inherited together because recombination is suppressed (Thompson and Jiggins, 2014, Berdan et al., 2022b, Schwander et al., 2014). In fact, sex chromosomes are generally considered a kind of supergene (Schwander et al., 2014). Supergenes come in (at least) two versions and which version(s) an individual possesses – two different versions or the same version twice – may greatly influence its phenotype. Hence, zygosity is of great biological relevance and techniques that allow zygosity to be determined would be widely applicable, given the widespread occurrence of sex chromosomes and other supergenes across the tree of life. Ideally, such a technique would be highly capable of processing many samples simultaneously and rapidly at low cost.

Kompetitive Allele Specific PCR (KASP) is an efficient and cost-effective method for genotyping SNPs in a large number of samples (Semagn et al., 2013). In this method, typically a single common reverse primer is used in combination with two allele-specific forward primers that differ in the last nucleotide, defined by the targeted SNP. These primers also possess a tail that is complementary to one of the two quenched fluor-labelled oligos; one with FAM, one with HEX. When thermal cycling exponentially increases one tail (in individuals homozygous for one or the other SNP) or both tails (in heterozygous individuals), the complementary label is no longer quenched, resulting in a corresponding fluorescent signal. However, with a slight modification, KASP could, in theory, also be used to efficiently genotype markers for zygosity. Rather than targeting a single marker with SNP-specific primers, as in ‘classical’ KASP, two different (i.e. non-homologous) markers that are positioned on the genomic region relevant for zygosity could be targeted, to determine the presence or absence of these markers. To this aim we use two distinct primer pairs, each with a different quenched fluor-labelled oligo, in a single reaction (Fig. 1). We call this modification multiplex KASP (mxKASP).

**Figure 1.**
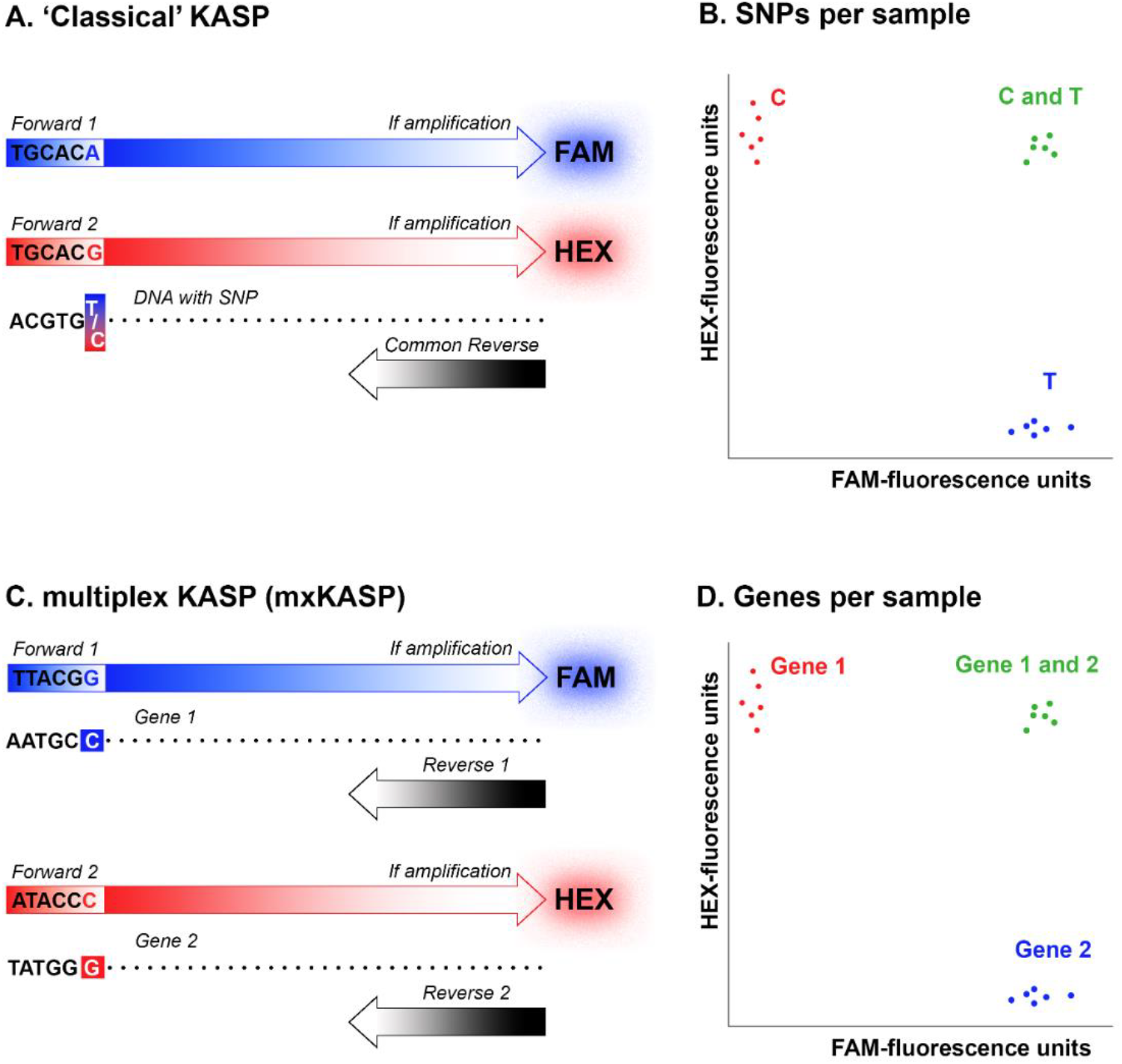
Schematic showing the different approach between KASP and mxKASP. (A) In KASP two forward primers differ in their last nucleotide as defined by the targeted SNP. Depending on which SNP variant is present, amplification leads to a fluorescent signal of FAM, HEX, or both. (B) For individuals homozygous for one SNP variant (in this case C) only the HEX signal is amplified (red dots), while for individuals homozygous for the other SNP variant (T) only the FAM signal is amplified (blue dots); for heterozygous individuals (C/T) both fluorescent signals are emitted (green dots). (C) In mxKASP two pairs of forward and reverse primers are designed for different (non-homologus) genes. (D) The presence of only one or the other gene will result in either the HEX (in this case for gene 1, red dots) or the FAM (gene 2, blue dots) signal to be amplified, while if both genes are present, both fluorescent signals are emitted (green dots).

We show proof of concept by applying mxKASP in a balanced lethal system. In a balanced lethal system two versions of a chromosome are involved (Berdan et al., 2022a, Wielstra, 2020, Muller, 1918). Healthy individuals possess a single copy of both chromosome versions, meaning they are reciprocally hemizygous, while diseased individuals possess two identical chromosome versions, meaning they are homozygous for one or the other version. The best studied example of a balanced lethal system concerns the *chromosome 1 syndrome* in newts of the genus *Triturus*, in which two versions of chromosome 1 are involved: 1A and 1B (Sessions et al., 1988, Sims et al., 1984, Macgregor and Horner, 1980). Only individuals that possess both versions of chromosome 1 (1A1B) survive embryonic development, whereas individuals that have the same version twice (1A1A or 1B1B), and therefore lack the alternate version, perish inside the egg. We design KASP markers that are positioned either on 1A or 1B, with the expectation that only one or the other marker would be amplified in diseased (1A1A or 1B1B) individuals, whereas both should amplify in healthy (1A1B) individuals.

## Material and Methods

### Sample selection

Embryos representing nine *Triturus* species (Wielstra and Arntzen, 2016), including the two distinct Balkan and Italian clades of *T. carnifex* (Wielstra et al., 2021) were included in the study. We included three individuals per genotype (1A1B, 1A1A, and 1B1B), adding up to 90 *Triturus* individuals in total. We also included three individuals of *Lissotriton vulgaris*, a species that is closely related to *Triturus* (Rancilhac et al., 2021), but that is not affected by the balanced lethal system (Horner and Macgregor, 1985).

### Marker selection

(De Visser et al., 2024b) produced DNA sequence data for c. 7k nuclear DNA markers from across the genome of three healthy (1A1B) and six diseased (three 1A1A and three 1B1B) individuals for the same ten *Triturus* clades as above (i.e. 90 samples in total). This dataset includes dozens of 1A-linked markers (not present in 1B1B individuals) and 1B-linked markers (not present in 1A1A individuals), next to thousands of additional markers positioned elsewhere on the genome. We also included three individuals of *Lissotriton vulgaris* from (De Visser et al., 2024a). We took sequences of the (in *Triturus*) 1A-linked marker *PLEKHM1*, the 1B-linked marker *NAGLU* and the ‘control marker’ *CDK17*, found on another linkage group than the 1A and 1B marker (De Visser et al., 2024c, France et al., 2024). Sequences were aligned and trimmed in MEGA 11 (Tamura et al., 2021), using the Muscle alignment tool set. Next, Unipro UGENE v.33 (Okonechnikov et al., 2012) was used to generate consensus sequences, with polymorphisms encoded by the relevant IUPAC codes. The consensus sequences were generated using an 80% threshold value.

### mxKASP

Four primer pairs were designed for the 1A-linked marker *PLEKHM1* and an additional four for the 1B-linked marker *NAGLU*, using LGC genomics’ Kraken v.23.11. This resulted in 16 (four by four) possible primer pair combinations (Supplementary Table S1, see Zenodo). KASP assays were prepared in a total volume of 200 μl, with 24 μl of both forward primers (10 μM), 60 μl of both reverse primers (10 μM), and 32 μl purified water. Next, KASP mix was made for each primer combination and contained 1.26 μl KASP assay, 102.8 μl KASP master mix and 102.8 μl purified water. All 16 combinations were tested in a 1,536 well plate with 1 μl of KASP mix and 1 μl of DNA for 93 samples (90 *Triturus*, 3 *Lissotriton*), as well as a negative control (no DNA). mxKASP was performed in a hydrocycler, beginning with 10 cycles with an annealing temperature decreasing from 61 to 55°C, followed by 26 cycles at 55°C. Two additional three-cycle recyclings were performed with annealing at 57°C. Fluorescence was measured using a PHERAstar plate reader and data were analysed using Kraken.

Two primer pair combinations (1A-linked primer pairs NAGLU.2 or NAGLU.3 with 1B- linked marker primer pair PLEKHM1.3; Table 1) showed separation into three clear groups (see results), corresponding to the three potential genotypes (1A1A, 1A1B, 1B1B). These two combinations were repeated in a 384 well plate to confirm the results. The KASP mix for each primer pair consisted of 7.2 μl assay, 194.4 μl KASP master mix and 194.4 μl purified water. Here, 3 μl of DNA of the same samples, next to a negative control, were combined with 3 μl of KASP mix, with identical downstream procedures as before.

**Table 1.**
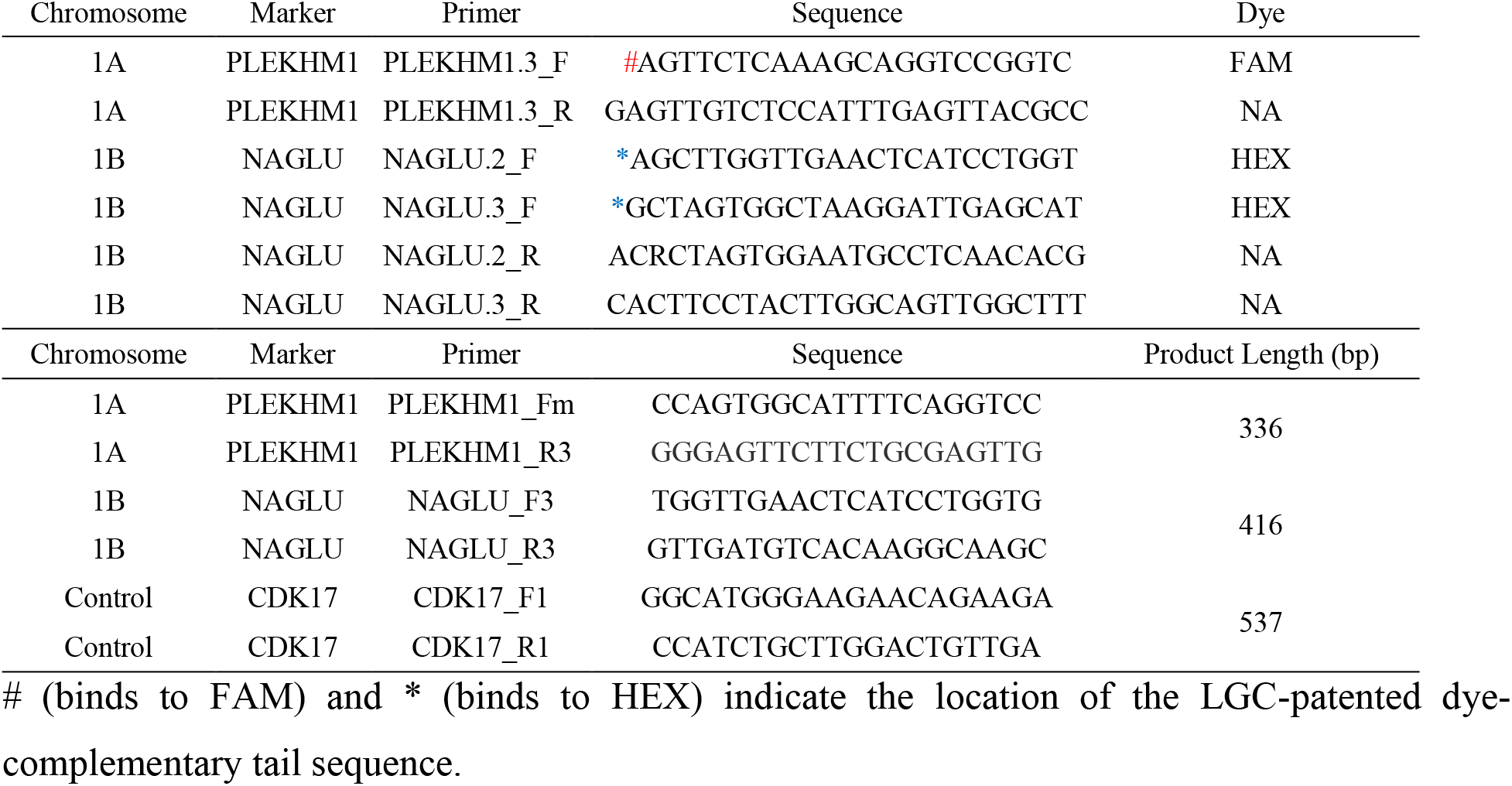
List of primers used for mxKASP (top six) and mxPCR (bottom six).

### mxPCR

To confirm our mxKASP results we designed a multiplex PCR (mxPCR) protocol. Primers were developed in Primer3Plus (Untergasser et al., 2012), aiming for a size difference (to be able to distinguish markers by gel electrophoresis) between the 1A-linked marker *PLEKHM1* (200- 300bp), the 1B-linked marker *NAGLU* (350-450bp) and the control marker CDK17 (>500bp). All other parameters were kept the same (Table 1). The mxPCR was performed using QIAgen multiplex PCR master mix. PCR was performed in 12 μl reactions, containing 0.06 μl of all forward and reverse primers (at 10 μM concentration, resulting in a 0.05 μM end concentration of each primer), 6 μl QIAGEN multiplex PCR master mix, 4.64 μl purified water and 1 μl of DNA. PCR conditions were as follows; a hot start for 15 minutes at 95°C, followed by 35 cycles of denaturation for 30 seconds at 95°C, annealing for 1 minute at 56°C, and extension for 1 minute at 72°C, and a final ten-minute extension at 72°C.

## Results

In our mxKASP experiment, two out of sixteen combinations of primer pairs resulted in a clear separation of three genotype clusters of the samples (Fig. 2; Supplementary Table S2, see Zenodo). The combination PLEKHM1.3 and NAGLU.2 showed allocation of all individuals into the expected genotype clusters, while two 1A1B samples were outliers for the combination of primers PLEKHM1.3 and NAGLU.3 (Fig. 2). The mxPCR amplification genotyped each individual as expected (Fig. 3).

**Figure 2.**
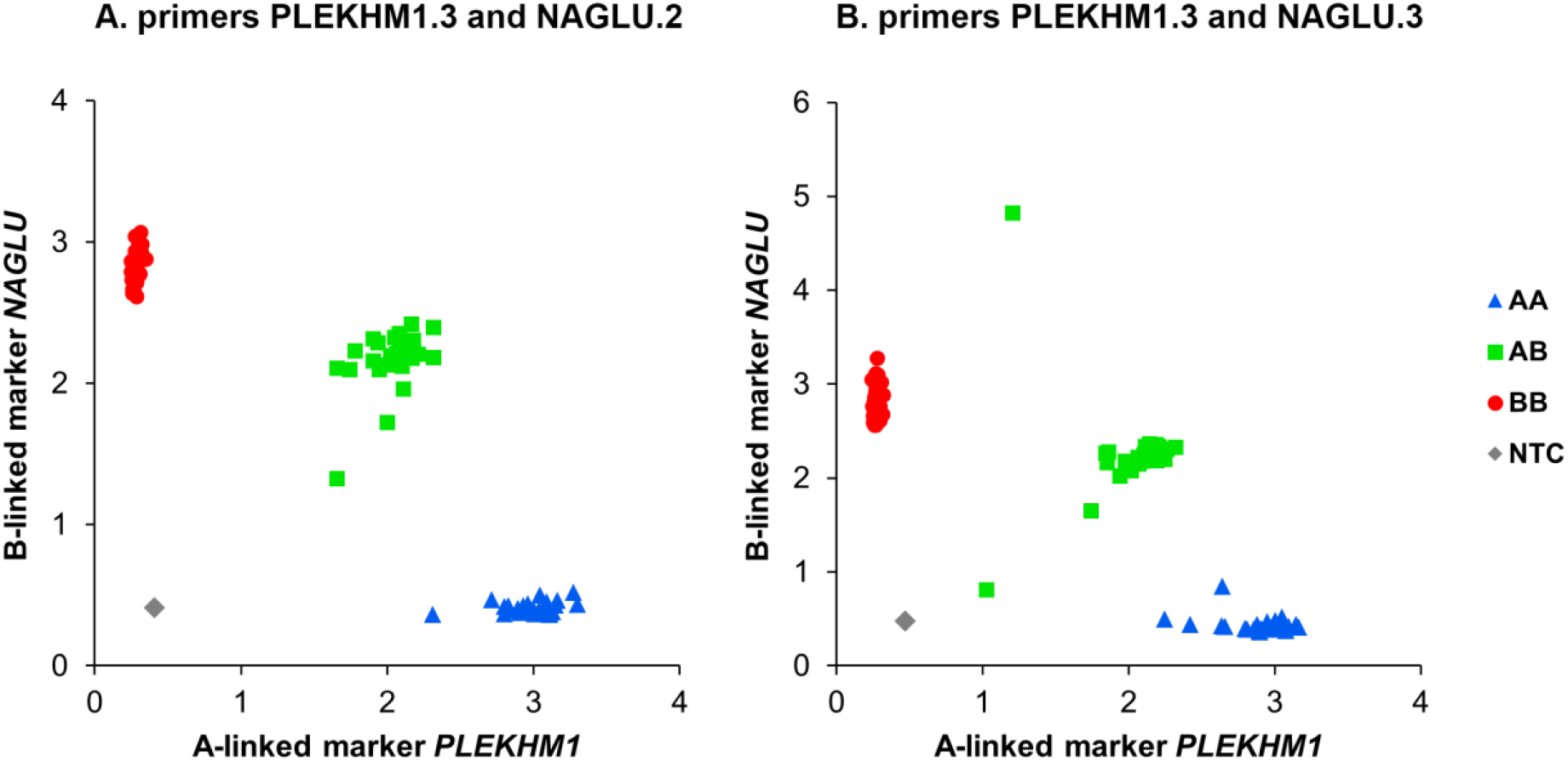
KASP results for two markers on different chromosomes, one on chromosome 1A (*PLEKHM1*) and one on chromosome 1B (*NAGLU*). Primers PLEKHM1.3 and NAGLU.2 (A. left panel) lead to complete separation of the samples into three clusters, while primers PLEKHM1.3 and NAGLU.3 (B. right panel) show a similar division with outliers for two 1A1B individuals. The x-axis reflects FAM and the y-axis HEX fluorescence units (y-axes are not to scale).

**Figure 3.**
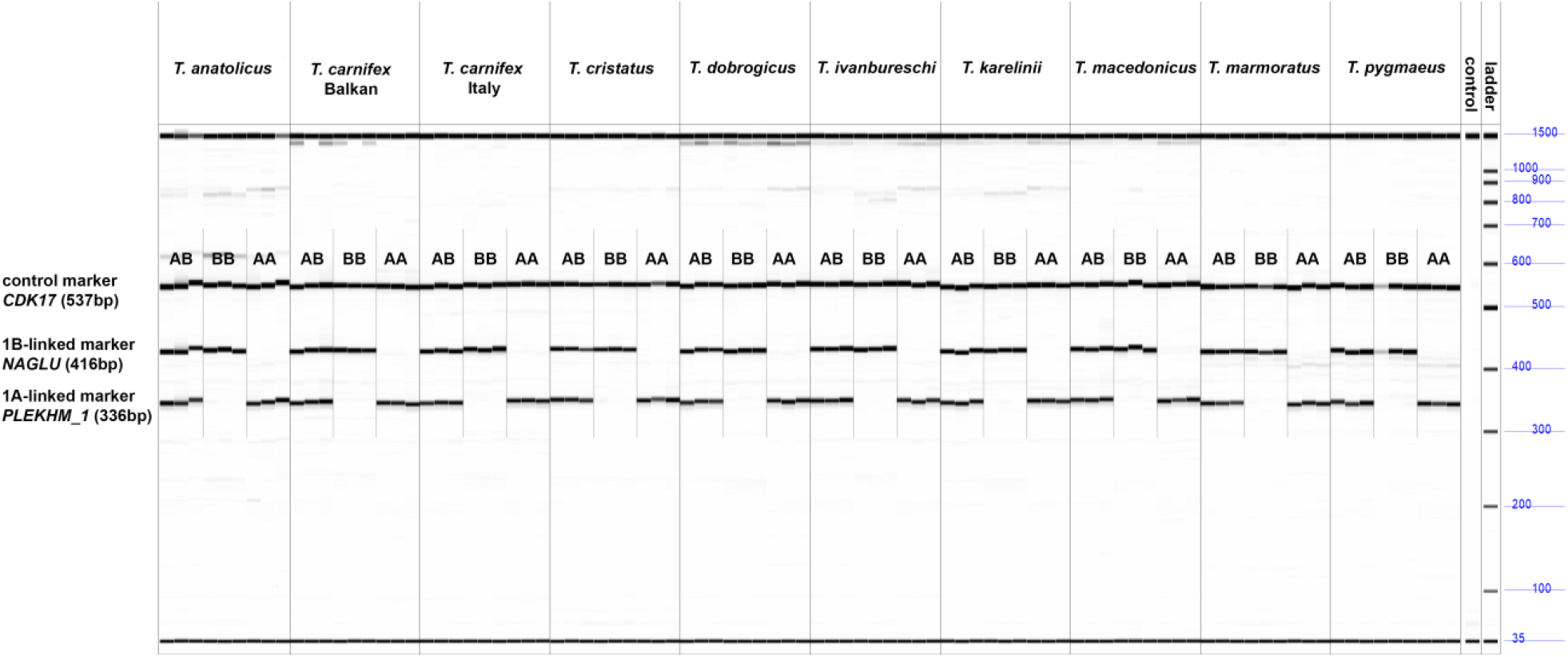
Multiplex PCR results obtained with a Fragment Analyzer. Three genotypes (healthy 1A1B or diseased 1A1A or 1B1B) can be distinguished for ten *Triturus* (candidate) species.

## Discussion

Our modification of KASP, mxKASP, proved to be successful in determining zygosity. Instead of targeting different SNPs on a single marker, as is typically done with KASP, we show that it is possible to target completely different, non-homologous markers. We use mxKASP to independently establish the presence or absence of two markers, which allows us to infer zygosity in the *Triturus* balanced lethal system. ‘Classical’ KASP has been used before to determine zygosity by targeting a SNP distinguishing the X and Y chromosomes, that could be identified because it co-segregated with the male-specific region (She et al., 2021). Our approach does not require such a SNP discovery step; rather it would directly target a marker private to the male-specific region, next to a marker present in both males and females.

Earlier KASP modifications could potentially inform on zygosity. ‘Duplex KASP (dKASP)’ was shown to be able to amplify two SNPs on two paralogous genes in a single run, with one of the two potential SNP variants for each paralog being present, resulting in a fluorescent signal (Jiang et al., 2023). This could be applicable to non-homologous genes as well, but depends on the presence of SNPs, which is not the case for mxKASP. The KASP modification ‘4- fluorescent KASP’ amplifies two different SNPs, positioned on two non-homologous markers, in a single run, by using four rather than the standard two fluorescent cassettes (Suo et al., 2020). Our mxKASP approach is simpler because it does not require additional fluorescent cassettes and is not dependent on the presence of SNPs on the markers of interest.

Over the last two decades KASP has become increasingly popular as a uniplex SNP genotyping platform for the agricultural, biological, forensic, and medical sciences (Kalendar et al., 2022, Majeed et al., 2018, Kaur et al., 2020). There are several reasons as to why it is applied so broadly. Firstly, KASP is capable of processing large sample sizes: it can be run in 96, 384 or 1,536 well-plates, with up to fourteen plates per PCR program (Semagn et al., 2013). Secondly, genotyping data interpretation in Kraken is practically automated, as well as instantaneous. In comparison, the capacity of general uni- or multiplex PCR approaches is limited by the number of available PCR machines and data interpretation is relatively time consuming, relying on labor intensive gel or capillary electrophoresis. Therefore, when sample sizes are large, KASP (including modifications such as mxKASP) is more efficient.

Acquiring clean-cut KASP results depends – as is the case for any genotyping method – on the quality of input DNA (Lear et al., 2018, Cavanaugh and Bathrick, 2018, Brusa et al., 2021, Hsiao, 2019). However, KASP only requires low quantity and quality DNA. Therefore, if KASP is combined with quick and cheap upstream methods, this opens up possibilities for scientists to upscale their research to the point of being able to sample, extract and genotype thousands of individuals in a matter of days (Wielstra et al., 2016). For this reason, KASP has already been embraced as a more economical choice for quality control analysis in for instance plant breeding and seed systems, rather than genotyping by sequencing (Ertiro et al., 2015, Semagn et al., 2013). The same applies to the clinical and medical fields, which incorporate KASP to rapidly and accurately identify blood groups for blood donors (Krog et al., 2019), as well as for screening for genetic susceptibility to diseases such as Parkinson’s and thrombosis (Altwayan et al., 2023, Landoulsi et al., 2017).

We foresee multiple applications in which our KASP modification, mxKASP, can be used for large-scale zygosity level determination. The method can be used to improve animal welfare. For instance, mxKASP could replace capillary electrophoresis of PCR products or real-time quantitative PCR previously recommended for use in the poultry industry to sex embryos *in ovo* to reduce male chick culling (He et al., 2019). Any PCR-based method would avoid the controversial approach of genetically modifying layer hens for determining the sex of their eggs (Bruijnis et al., 2015). mxKASP could also prove valuable in a conservation context, for example, in species where sex is influenced by both a genetic and a thermal component (Hattori et al., 2012, Geffroy and Wedekind, 2020). Circa 1/3 of fish species thought to exhibit only temperature-dependent sex determination appears to possess genetic sex determination too, and which one predominates depends on environmental factors (Ospina-Alvarez and Piferrer, 2008). Global warming threatens such species by biasing sex ratios. As more sex-linked markers are discovered, more genotype/phenotype mismatches will presumably be revealed (Edmands, 2021, Geffroy and Wedekind, 2020). Finally, mxKASP can be used to detect zygosity in fundamental eco-evolutionary studies of supergene systems. For example, to test the ratios of different genotypes when a particular supergene version is assumed to carry recessive deleterious mutations (Kupper et al., 2016, Tuttle et al., 2016).

## Author contributions

W.R.M.M., M.C.dV. and B.W. conceived the research. W.R.M.M., M.C.dV and A.T. conducted labwork. W.R.M.M. and M.C.dV analyzed the data. M.F. provided samples. W.R.M.M., M.C.dV. and B.W. wrote the manuscript with input from the other authors.

## Acknowledgements

This project received funding from the Dutch Research Council (NWO Promotiebeurs voor leraren 023.016.006) and the European Research Council (ERC) under the European Union’s Horizon 2020 research and innovation programme (Grant Agreement No. 802759). Salamander embryos were obtained by MF from his personal breeding colony. Housing and breeding salamanders as a private individual are not considered to require a license (BGBl. I S. 3125, 3126, 3750). Housing and breeding protocols comply to EU directive standards (EU directive annex III, section B, Table 9.1) and were reviewed by the Animal Welfare Body Leiden. As per EU legislation regarding the protection of animals used for scientific purposes (EU directive no. 2010/63/EU), sacrificing embryos that are not feeding independently does not qualify as an animal experiment. Carola Feijt, James France, Bobbie Sewalt and Klaas Vrieling helped with laboratory work.

